# GrAdaBeam: Combining model gradients with evolutionary search for generalizable nucleic acid design

**DOI:** 10.1101/2025.06.20.660785

**Authors:** Joel Shor, Erik Strand, Cory Y. McLean

**Affiliations:** Move37 Labs, Somerville, MA, USA; MIT Center for Bits and Atoms, MIT, 77 Massachusetts Ave, Cambridge, 02139, MA, USA; Google, 355 Main Street, Cambridge, 02142, MA, USA

**Keywords:** nucleic acid, generative design, therapeutics

## Abstract

We introduce GrAdaBeam, a hybrid model-based optimization algorithm that combines gradient-derived attention maps with an adaptive beam search to navigate complex nucleic acid fitness landscapes. By unifying the broad exploration of evolutionary methods with the precise guidance of gradient descent, GrAdaBeam overcomes a central limitation of existing approaches: no single optimization strategy performs robustly across the full spectrum of genomic design tasks. We rigorously evaluate GrAdaBeam and seven other design algorithms using NucleoBench, a novel benchmark covering 17 diverse genomic tasks that introduces a paired-start-sequence design for superior statistical comparisons. GrAdaBeam statistically outperforms all other algorithms across over 600,000 experiments, never ranking lower than second across all 17 benchmark tasks, while baseline methods often struggle on large models or long sequences. Critically, GrAdaBeam sequences generalize most reliably to independent predictive models and recover canonical transcription factor binding motifs de novo, providing evidence of biological signal capture beyond the optimization target. GrAdaBeam and NucleoBench are freely available as an open-source package.

## 1 Introduction

Designing nucleic acid sequences with defined regulatory functions has many applications in precision medicine [1–5], including designing more precise CRISPR guide RNAs with minimal off-target effects for gene editing therapies [6], creating mRNA vaccines with optimized stability and translational efficiency [7], designing antisense oligonucleotides that maximize target binding while minimizing immunogenicity [3, 8], and developing aptamers with enhanced specificity for diagnostic applications [9].

Deep learning models can now effectively predict many aspects of genomic function from sequence [10–17]. However, engineering synthetic sequences that function with superhuman efficacy *in vivo* remains challenging.

To engineer these sequences, the field has split into two distinct approaches: generative modeling and model-based optimization (Fig. 1a). Generative models, such as diffusion models [18], autoregressive transformers [19], and generative adversarial networks [20, 21], learn the probability distribution of natural sequences. In their unconditional form, they excel at producing biologically plausible candidates but prioritize mimicry over maximization. Hybrid approaches that augment generative models with reward signals or reinforcement learning can partially bridge this gap, but the optimization component in such methods faces the same landscape-traversal challenges addressed here. In contrast, model-based optimization [22], the focus of this work, treats design as a search problem (Fig. 1b). By inverting predictive oracles to ascend a fitness landscape, these methods seek “supernatural” sequences with efficacy beyond evolutionary limits, but sometimes run the risk of finding biologically-implausible solutions.

**Fig. 1.**
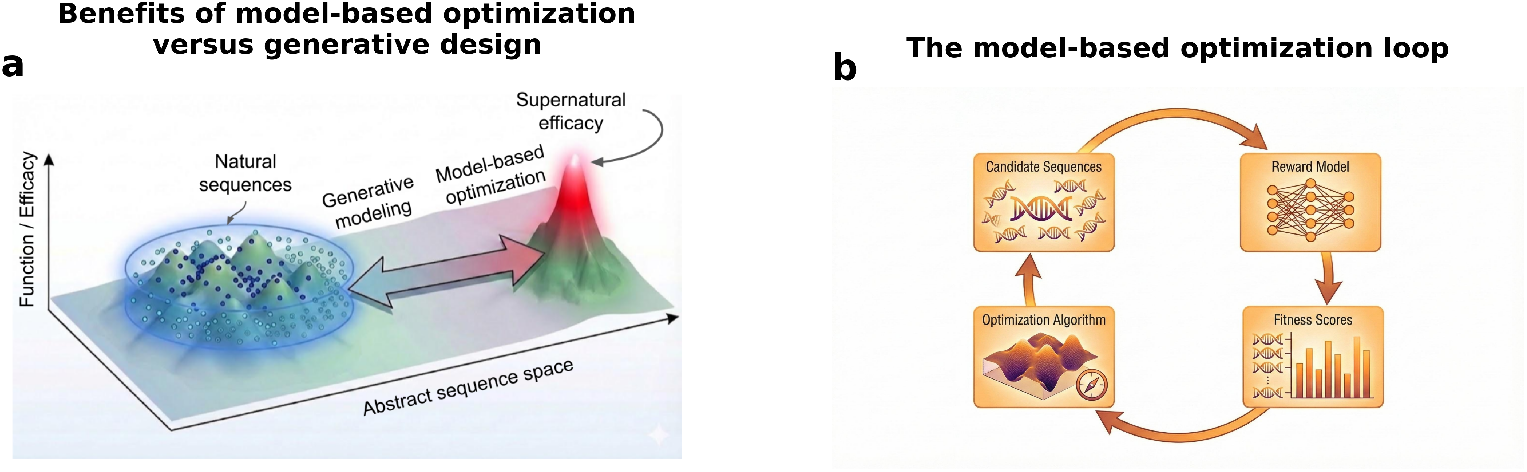
Model-based optimization prioritizes efficacy, while unconditional generative design prioritizes mimicry. **a)** Unconditional generative modeling learns a distribution from natural sequences and samples from it. Model-based optimization attempts to find the best sequence, regardless of how “natural” or “common” it is. Hybrid generative approaches exist but still require an optimization component to exceed the training distribution. **b)** Model-based optimization relies on an iterative process of running a model, using the results to determine new candidate sequences, and repeating.

Current computational designers using model-based optimization face the choice between evolutionary methods and gradient methods. While other optimization frameworks exist [23–25], evolutionary and gradient methods dominate nucleic acid design practice because they operate natively on discrete sequences without requiring surrogate models, continuous relaxations, or policy training. Evolutionary methods use random mutations to improve sequences and treat objectives as “black boxes,” while gradient methods use derivatives of the objective function to guide mutations. Each method succeeds on different tasks (Fig. 2), and both methods are capable of converging on high-scoring sequences that exploit mathematical artifacts in the model rather than biological reality. Consequently, *in silico* designs do not always reliably translate to biological function, a gap that remains difficult to close without rigorous independent validation.

**Fig. 2.**
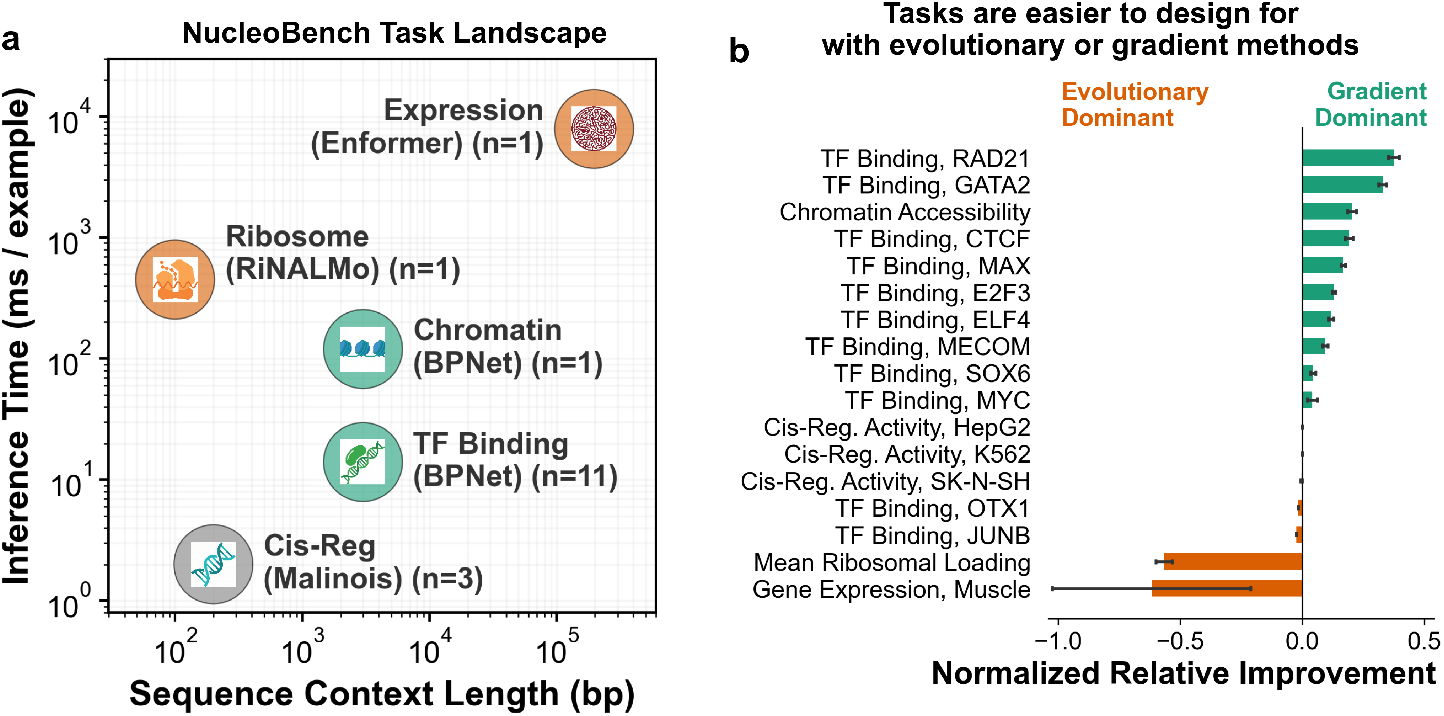
NucleoBench tasks and the performance divide between evolutionary and gradient designers. **a)** NucleoBench includes 17 biological tasks spanning five categories that vary multiple orders of magnitude across both sequence length and model inference speed. Evolutionary algorithms are the best performers on orange tasks, gradient algorithms are best on green tasks. **b)** Normalized relative improvement of gradient-based methods over evolutionary-based methods for each NucleoBench task. Positive values (green) denote superior performance by gradient-based methods and negative values (orange) denote superior performance by evolutionary methods. Error bars indicate the 95% confidence intervals derived from the standard errors of the means. Gray bars denote comparisons where the performance difference is not statistically significant (95% CI includes zero).

In this work, we present a unifying solution to the choice that computational designers face between evolutionary and gradient methods: we introduce a designer that integrates gradient guidance with adaptive sampling (GrAdaBeam), which traverses complex fitness landscapes, produces designs that generalize across independent biological models, and recovers native regulatory syntax.

## 2 Results

### 2.1 NucleoBench: An evaluation framework for model-based nucleic acid design algorithms

We created NucleoBench, a benchmark that enables a fair-comparison for testing model-based optimization of nucleic acid sequences. An “experiment” in NucleoBench consists of a task (ex. “binding affinity of the GATA2 transcription factor”), a designer (ex. “simulated annealing”), a collection of hyperparameters for that designer (ex. annealing rate=1.0, …), and a start sequence (ex. ATGC…). The design algorithm attempts to iteratively improve the sequence for a fixed amount of time. For more details on the NucleoBench framework, see Section 4.1.

NucleoBench consists of 17 tasks across five categories that span a range of sequence length, model complexity, and biological complexity (Fig. 2a, Table 1). The tasks are predicting **cis-regulatory activity** (*n* = 3), **transcription factor binding** (*n* = 11), **chromatin accessibility** (*n* = 1), **mean ribosomal loading** (*n* = 1), and **cell-type gene expression** (*n* = 1). For more details on the tasks, see Section 4.2.

**Table 1.** Summary of design tasks in NucleoBench. *Input length is 196,608 bp, but only 256 bp are edited.

| TASK CATEGORY | MODEL | NUM TASKS | SEQ LEN (BP) | INFERENCE (MS / EXAMPLE) | GRADIENTS (MS / EXAMPLE) |
| --- | --- | --- | --- | --- | --- |
| Cell-type specific Cis-regulatory activity | Malinois [14] | 3 | 200 | 2 | 6 |
| Transcription factor binding | BPNet-lite [12] | 11 | 3000 | 14 | 30 |
| Chromatin accessibility | BPNet-lite [12] | 1 | 3000 | 120 | 320 |
| Ribosomal loading | RiNALMo [13] | 1 | 100 | 450 | 1300 |
| Selective gene expression | Enformer [10] | 1 | 196,608 / 256 * | 7900 | 27000 |

We evaluated NucleoBench on eight designers. Four are leading model-based design methods in the literature, either by popularity or performance (directed evolution, simulated annealing, ledidi, adalead), two are algorithms from computer science optimization literature (unordered beam search, beam search), and two are algorithms introduced in this paper (gradient-guided directed evolution, gradient-guided adaptive beam search). The designers span a number of axes, with an important one being whether they leverage model gradients (*n* = 5 do not, *n* = 3 do). For more details on the designers, see Section 4.3.

Initial benchmarking of the baseline designers show that design performances of gradient versus non-gradient methods are highly complementary (Fig. 2b). This performance split exposes a fundamental limitation in current computational design: no single existing strategy is robust across the full spectrum of genomic tasks. Relying primarily on gradients risks failure in complex therapeutic contexts (e.g., mRNA stability), while pure evolutionary methods are computationally inefficient in other contexts (e.g. for finding local motifs). This complementary failure mode requires a hybrid architecture capable of dynamic blending between different approaches.

### 2.2 GrAdaBeam: A hybrid gradient-evolutionary design algorithm

To resolve the dichotomy between gradient and evolutionary exploration, we developed a simple hybrid algorithm “Gradient Evo” (Section 4.3.3), and “GrAdaBeam” (Gradient-Guided Adaptive Beam search, Fig. 3a). While Gradient Evo improved over pure evolutionary methods on gradient-favorable tasks, it lacked the beam structure and adaptive hyperparameter tuning needed for consistent cross-task performance, motivating the full GrAdaBeam architecture. GrAdaBeam and Gradient Evo constrain the vast search space using gradient-derived “attention maps.” Specifically, they utilize the predictive model’s gradients to compute a Taylor *in silico* mutagenesis (TISM, Eq. (2)) [26] map for every candidate, which is used to identify potentially high-impact nucleotides. GrAdaBeam and Gradient Evo oversample from these positions when generating mutations. GrAdaBeam uses a beam search [27] to prevent plateauing in local optima. GrAdaBeam includes dynamic blending between gradient signals and random mutations, enabling it to outperform existing designers regardless of which style the task is better suited to (Fig. 2b). It also enables something unique: **GrAdaBeam is able to take advantage of <u>parts</u> of the task landscape that are more easily traversed using gradient guidance**.

**Fig. 3.**
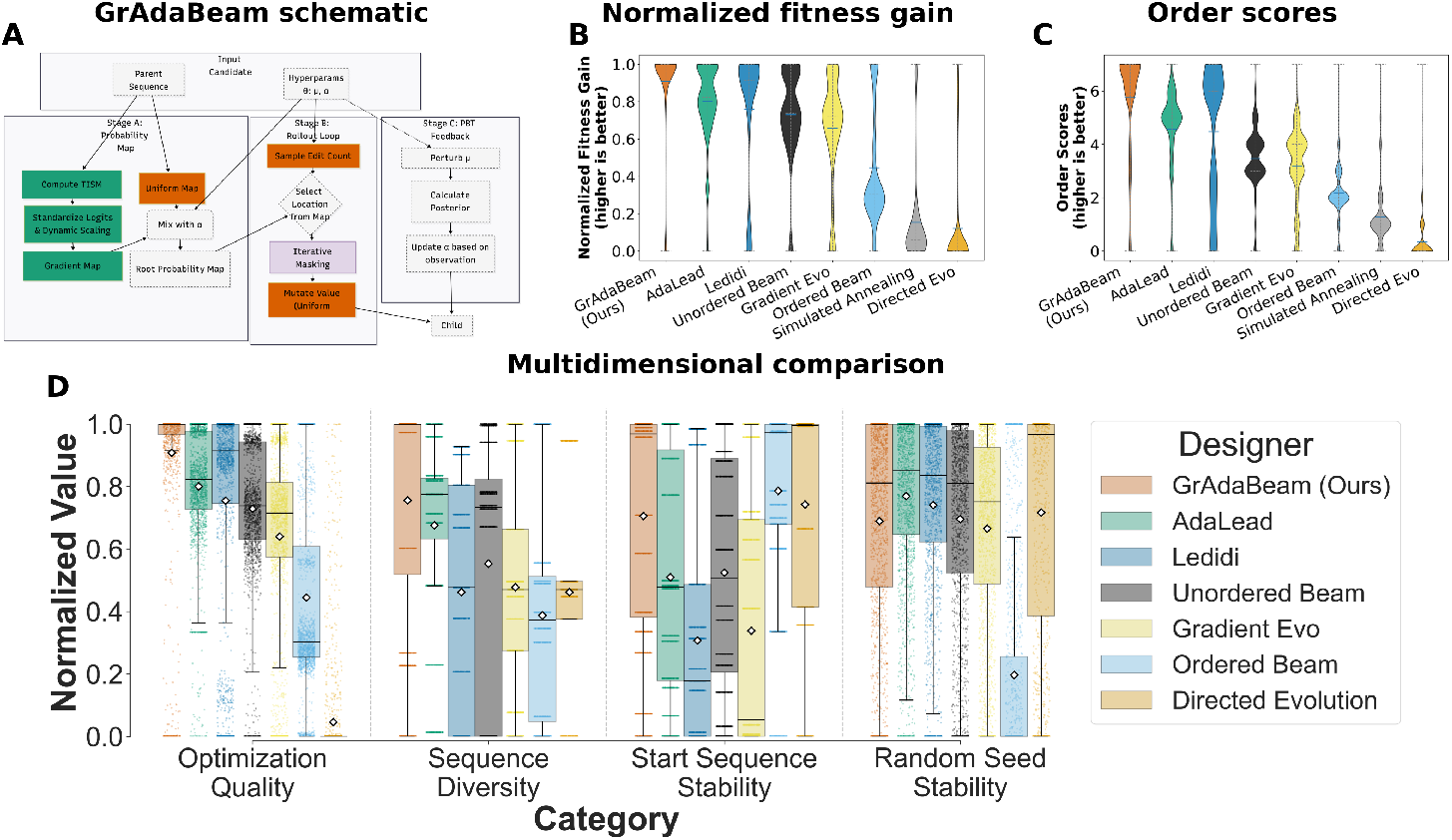
GrAdaBeam demonstrates superior performance, diversity, and robustness across 17 biological tasks. **a)** The GrAdaBeam algorithm decomposes optimization into three probabilistic steps, blending gradient exploitation with evolutionary exploration. Orange boxes represent evolutionary steps, green are gradient steps. See Section 2.2 for an overview of the method, and Section 4.3.1 for the details. b)Violin plots showing the distribution of normalized fitness improvements relative to the starting sequence. Raw gains are normalized by the standard deviation of the start sequences’ fitness and linearly rescaled to a [0, 1] interval to allow for direct comparison across tasks with varying ranges. 1.0 indicates the top-performing designer. C)Violin plots showing the rank-based reliability of each algorithm. For a given task and start sequence, the order score indicates the number of designers with worse final fitness. 7.0 indicates the top-performing designer. **d)** Box plots detailing algorithm performance across five dimensions: Optimization Quality (fitness), Performance on Orthogonal Models, Sequence Diversity (average pairwise Hamming distances), Start Sequence Stability (robustness across start sequences), and Random Seed Stability (robustness to random seeds). Values for all dimensions are standardized and normalized to a [0, 1] scale, where higher values indicate superior properties.

GrAdaBeam uses Population Based Training (PBT) [28] to adapt the strength of gradient-guidance versus a random walk. As the search progresses, the algorithm not only evolves the sequences but also evolves the search strategy itself: candidates that successfully improve fitness transmit their hyperparameters to the next generation, while failing strategies are discarded. This allows GrAdaBeam to autonomously transition from “high-exploration” (random mutations) to “high-exploitation” (gradient-driven refinement) as it traverses the fitness landscape. For the parameter governing the influence of the gradient attention maps, GrAdaBeam uses an adaptive parameter mechanism based on bayesian inference and sampling characteristics used to find the best sequences. See Section 4.3.1 for more details.

### 2.3 GrAdaBeam achieves superior fitness, diversity, and robustness across tasks

Benchmarking GrAdaBeam on 17 NucleoBench tasks against four established baselines and three novel controls revealed that GrAdaBeam provides a generalized solution to nucleic acid design. GrAdaBeam achieved the highest aggregate performance when measured both by sequence fitness gain (Fig. 3b) and the rank-based “Order Score” metric (Fig. 3c) (see Methods for details). Statistical analysis confirms this dominance: global pairwise Wilcoxon signed-rank tests with Holm-Bonferroni correction (*α* = 0.05) show that GrAdaBeam significantly outperforms all seven comparison algorithms across the benchmark suite (*p <* 0.002). On a per-task basis, GrAdaBeam ranks first in 9 of 17 tasks, and is statistically indistinguishable from the top-performing algorithm in 4 of the 17 tasks (AdaLead and Unordered Beam tied for first on all three cis-regulatory prediction tasks, unordered beam tied for best on gene expression prediction task). In the remaining four, it consistently achieves the second-best rank (outperformed by Ledidi on four transcription factor binding tasks). See Section 4.6 for details on the statistical test, and Supplementary Table 1 for task-specific, 95% confidence intervals across start sequences. Taken together, these results show that GrAdaBeam-designed sequences are consistently better than others for the *in silico* tasks that they were designed for.

Beyond raw score, a practical therapeutic designer must produce results that are diverse, reliable, and robust. We profiled the diversity of each designer’s set of outputs by measuring their average pairwise Hamming distance. We profiled each algorithm’s stability by measuring performance variance across different random seeds and start sequences. GrAdaBeam exhibited high diversity and stability (Fig. 3d). It consistently converged to high-fitness solutions regardless of the random seed initialization (Supplementary Table 2). Furthermore, it proved robust to start-sequence bias (Supplementary Table 3), effectively optimizing “difficult” start sequences that caused other methods to underperform (Supplementary Table 4). This combination of high fitness, population diversity, and algorithmic reliability establishes the method as a robust engine for discovery, capable of generating reproducible and varied candidates for experimental validation. See Section 4.5 for details on the metrics used to quantify these properties.

### 2.4 GrAdaBeam designs generalize across orthogonal predictive models

A central concern in model-based design is whether optimized sequences capture genuine biological properties or merely exploit characteristics of the oracle model. To assess whether GrAdaBeam recovers true biological signal rather than model-specific overfitting, we performed orthogonal validation on GrAdaBeam’s designed sequences by assessing their performance on independent, held-out predictive architectures and quantifying their recovery of native regulatory motifs (Fig. 4).

**Fig. 4.**
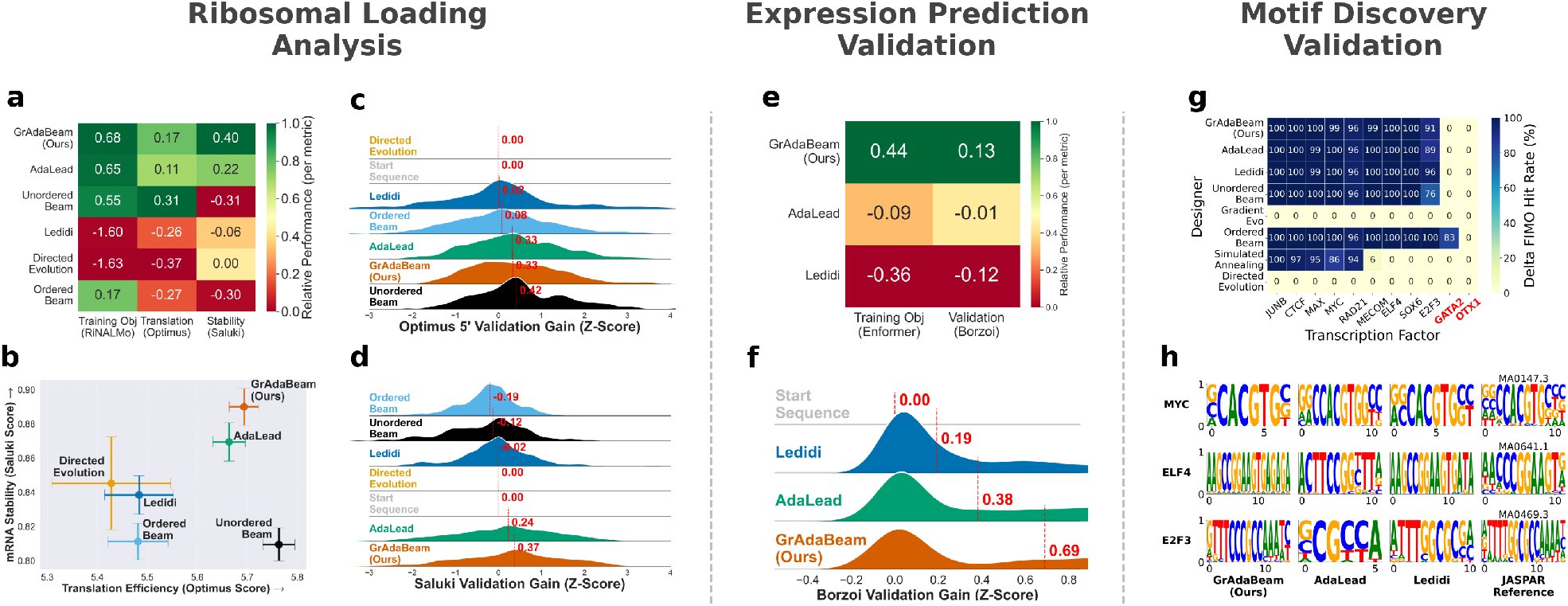
Orthogonal validation confirms designed sequences generalize across independent predictive architectures and recover native regulatory syntax. **a-d)** Sequences optimized for ribosomal loading on RiNALMo were evaluated using independent holdout models Optimus 5-Prime (translation efficiency) and Saluki (mRNA stability). **a**, Heatmap ranking design algorithms by a composite performance score. Color intensity is normalized column-wise (0–1) to display mean standardized performance gains. **b**, Trade-off map plotting mean absolute improvement in translation (x-axis) against mRNA stability (y-axis). GrAdaBeam sequences occupy the Pareto frontier (top-right quadrant), indicating concurrent optimization of both metrics. Error bars denote the 95% confidence interval. **c,d**, Ridgeline plot detailing the distribution of z-scored validation gains for Optimus 5’ (c) and Saluki (d). Distributions are modeled via Kernel Density Estimation (KDE) and ordered vertically by median gain (indicated by red dashed lines). **e, f)** Sequences optimized for cell-type-specific gene expression using Enformer were evaluated on the held-out Borzoi model. **e**, Heatmap displaying mean normalized performance, ranked by composite score. **f**, Ridgeline plot showing the distribution of z-scored Borzoi validation gains across algorithms. **g, h)** Validation of motif recovery in designed sequences. **g**, Heatmap of Delta FIMO Hit Rates, showing the increase in motif occurrence frequency in designed sequences relative to starting seeds. Rows represent design algorithms and columns represent target transcription factors (TFs), ordered by mean performance. **h**, Comparison of de novo motifs discovered in designed sequences (using STREME) against canonical JASPAR reference motifs for MYC, ELF4, and E2F3. All discovered motifs significantly match their respective reference PWMs (TOMTOM *p <* 5 × 10^*−*4^).

#### GrAdaBeam generalizes to held-out models of mRNA translation efficiency and stability

We performed orthogonal validation on the Ribosomal Loading task. While sequences were optimized using RiNALMo [13] (a model of ribosomal loading), we evaluated their quality using two independent, held-out architectures: Optimus 5’ [15] (which predicts translation efficiency on synthetic sequences) and Saluki [17] (which predicts mRNA half-life/stability on endogenous sequences).

The results demonstrate that GrAdaBeam does not merely exploit the source model, but optimizes underlying biological properties that generalize across architectures. As shown in Fig. 4a, GrAdaBeam achieved the highest aggregate validation scores, significantly outperforming pure gradient (Ledidi) and evolutionary (AdaLead) baselines on both held-out models. The distribution of validation gains (Fig. 4c,d) reveals that GrAdaBeam shifts the entire population of designed sequences towards higher biological fitness, whereas baseline methods often leave a substantial portion of candidates no better than the random starting sequences.

Most notably, our analysis uncovered that GrAdaBeam successfully navigates the complex trade-offs inherent in mRNA design. Fig. 4b maps this multi-objective landscape, plotting the mean improvement in translation (x-axis) against stability (y-axis). While standard methods like Directed Evolution struggled to improve both metrics simultaneously, GrAdaBeam sequences occupy the Pareto frontier (top-right quadrant), demonstrating statistically significant concurrent gains in both stability and translational output. This cross-model generalization strongly suggests that the algorithm has converged on genuine regulatory syntax rather than model-specific overfitting.

#### GrAdaBeam generalizes to held-out model of cell-type-specific gene expression

We performed orthogonal validation by assessing the designed sequences with an independent, held-out predictive model. The sequences in this task were optimized using the Enformer model [10] as the oracle. For validation, we used Borzoi [16], a distinct model also trained to predict CAGE-seq gene expression profiles from DNA sequence but featuring a different architecture and training data subset.

To enable direct comparison across models with different output distributions, raw prediction scores were converted to Z-scores normalized to the distribution of the start sequences. For a given model *M* (Enformer or Borzoi), a start sequence *s*_0_, and an optimized sequence *s*^*′*^, the optimization gain Δ*Z*_*M*_ is defined as:

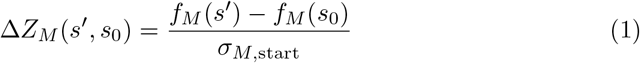

where *f*_*M*_(·) denotes the raw activity prediction of the model, and *σ*_*M*,start_ is the standard deviation of the raw scores across the pool of 100 start sequences. Note that the mean *µ*_*M*,start_ cancels out when calculating the difference. We utilized Spearman rank correlation to assess the consistency of optimization trends between the oracle and validation models. Additionally, we employed a one-sided Wilcoxon signed-rank test to determine if the designed sequences exhibited a statistically significant increase in predicted activity on the held-out model compared to the baseline. We present this relationship in Fig. 4e,f, plotting the optimization gain (Enformer) against the validation gain (Borzoi) for each sequence. As shown in the distributions, GrAdaBeam achieves the highest median validation gain, demonstrating robust generalization to the held-out Borzoi model compared to baseline methods.

#### Designed sequences recover known transcription factor binding motifs

To validate that the designed sequences rely on biologically relevant features rather than model artifacts, we assessed the recovery of known transcription factor (TF) binding motifs. We used FIMO [29] to scan both starting and designed sequences for the canonical JASPAR motifs corresponding to each target TF. Fig. 4g presents the “Delta FIMO Hit Rate,” defined as the increase in the percentage of sequences containing a significant motif match (*q <* 0.1) after optimization. This metric quantifies the extent to which each design algorithm actively inserts or preserves the correct regulatory syntax. We observed robust motif enrichment across most targets, with GrAdaBeam and AdaLead consistently achieving high recovery rates. Interestingly, the GATA2 and OTX1 motifs were challenging for all methods, potentially due to complex co-factor requirements or failures of the predictive model itself.

We further validated the fidelity of these inserted patterns by performing *de novo* motif discovery on the designed sequences using STREME [30] and aligning the discovered position weight matrices (PWMs) to the JASPAR database [31]. Visual inspection of the recovered motifs for representative TFs (MYC, ELF4, E2F3) reveals striking similarity to the canonical reference motifs (Fig. 4h). This structural similarity was confirmed quantitatively using TOMTOM [32], which aligned the discovered motifs to their JASPAR targets with high statistical significance (all comparisons *p <* 5 × 10^*−*4^). Notably, the MYC motif discovered in GrAdaBeam sequences matched the canonical MAX-binding E-box motif with exceptional precision (*p <* 10^*−*5^), suggesting the model has learned the precise biophysical constraints of the factor’s DNA binding domain.

## 3 Discussion

The transition from nucleic acid property prediction to generation has huge potential for medicine, from optimizing mRNA vaccines to designing precise gene therapies. However, this shift is currently hindered by two problems: existing design algorithms fail to traverse the optimization landscape of certain biological design problems, and designed sequences can have high *in silico* fitness but perform poorly in biological reality. We introduce the GrAdaBeam design algorithm to address both tensions.

By integrating the broad exploration of evolutionary algorithms with the precise guidance of gradient descent, GrAdaBeam outperforms state-of-the-art methods in 9 of 17 diverse genomic tasks, never ranks lower than second, and is the overall best designer by numerous metrics. In addition to finding better solutions than other designers and finding good solutions more consistently, GrAdaBeam exhibits high stability across start sequences and random seeds, and generates diverse candidate pools, which are practical requirements for use.

Importantly, our orthogonal validation results provide encouraging evidence that GrAdaBeam’s high-scoring sequences capture biological properties that extend beyond the specific oracle used during optimization. A key concern in model-based design is that optimizers can identify shortcuts specific to the oracle model rather than genuine regulatory syntax, and cross-model generalization is one meaningful indicator that this risk is reduced. We addressed this through rigorous orthogonal validation, using unsupervised sequence enrichment tools to compare biological motifs and held-out models with distinct architectures. GrAdaBeam-designed sequences optimized on RiNALMo (for ribosomal loading) showed significant performance gains when evaluated on Optimus 5’ (translation efficiency) and Saluki (mRNA stability). Similarly, sequences optimized for Enformer generalized robustly to Borzoi. This cross-model generalization, combined with the successful de novo recovery of known JASPAR transcription factor binding motifs, provides encouraging evidence that the algorithm is capturing true biological signals rather than model artifacts, although wet-lab validation remains necessary to confirm this.

Methodologically, GrAdaBeam advances the field of model-based optimization design of nucleic acids in multiple ways. First, it alleviates the need to choose evolutionary or gradient methods by blending the two together. GrAdaBeam uses Populated Based Training (PBT) and a Bayesian, adaptive parameter mechanism to allow the design landscape to dictate whether gradients or random mutations are emphasized. Second, the trend toward larger predictive models had previously prevented the most successful model-based designers from using the latest models. GrAdaBeam’s computational efficiencies of amortizing gradient computations (“iterative masking”) and O(1) sampling allows model-based designers to use the latest predictive models to design sequences. These same efficiencies also allow designers to create longer sequences.

The introduction of NucleoBench provides the necessary framework to rigorously measure these advancements. Prior to this work, the lack of standardized tasks and metrics fragmented the field, obscuring whether new methods offered genuine improvements or merely selected advantageous baselines. By evaluating designers across a standardized suite of 17 tasks, spanning a wide range of sequence lengths, biological complexities, and model architectures, NucleoBench exposes the limitations of relying on any single evaluation strategy. Crucially, its paired start sequence design enables robust, non-parametric statistical comparisons. For therapeutic applications, this comprehensive evaluation is vital. Our benchmarking framework uniquely highlights this, showing that sequences designed by GrAdaBeam consistently occupy the optimal quadrant of the trade-off map.

Our study has limitations that merit discussion. First, while orthogonal validation on held-out models significantly reduces the likelihood of hallucination, it is not a substitute for wet-lab verification. The sequences generated here are optimized *in silico*; their ultimate biological fidelity depends on the accuracy of the underlying oracle models (Malinois, BPNet, RiNALMo, Enformer). If these models contain systematic biases, our optimizer will inevitably inherit them. Directly incorporating model uncertainty is one direction that could address this. Second, future work must address the integration of explicit biological constraints, such as immunogenicity filters or secondary structure requirements, directly into the search loop to further bridge the gap between computational design and clinical application. Third, this work compared model-based optimizers, since those prioritize superhuman efficacy over mimicry. Recent generative approaches that incorporate reward-guided fine-tuning or reinforcement learning blur this boundary, and expanding NucleoBench to include such hybrid generative methods would give practitioners a more complete understanding of the design landscape.

Ultimately, the goal of nucleic acid design is to move beyond the discovery of natural variations to the engineering of synthetic sequences with superhuman efficacy. By resolving the dichotomy between gradient and evolutionary search, and by establishing a rigorous benchmark for validation, this work offers a robust foundation for that transition. We anticipate that the principles established here, such as dynamic search adaptation, orthogonal validation, and standardized benchmarking, will spur the development of next-generation algorithms capable of unlocking the full therapeutic potential of the non-coding genome.

## 4 Methods

### 4.1 The NucleoBench Evaluation Framework

To rigorously evaluate nucleic acid design algorithms, we developed NucleoBench, a standardized framework comprising 17 diverse genomic tasks. Each experimental run is defined by a tuple (*D, T, H, S*_0_), where *D* is the design algorithm, *T* is the design task (oracle model), *H* are the hyperparameters, and *S*_0_ is the starting sequence. Over 600,000 experiments were conducted on Google Cloud Platform using <monospace>n1-highmem-16</monospace> CPU instances. All experiments were capped at a fixed wall-clock time (12 hours for Enformer tasks, 8 hours for all others) rather than step count. We discuss the designers *D* in Section 4.3, tasks *T* in Section 4.2.

The **hyperparameters** are designer-specific. These are factors such as “learning rate” in Ledidi, and “mutation rate” in Directed Evolution. We used hyperparameter values that perform well in other benchmarks [14, 33–35], as well as adaptations (e.g. lower mutation on problems with longer sequences). More details on the number of hyperparameters used for each designer can be found in Supplementary Section “Number of hyperparameters”.

To ensure statistical power and disentangle algorithm performance from initialization bias, we employ a **paired start sequence design**. For every task *T*, we generated a fixed set of *N* = 100 start sequences 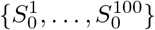. Every algorithm *D* is evaluated on this identical set. This paired design enables the use of more appropriate comparisons of performance (comparing designers from the same start sequence), orthogonal validation (comparing designed sequences from the same start sequence on hold-out models), and sensitive non-parametric statistical tests (Friedman test [36] with Nemenyi post-hoc) to identify performance differences and “intrinsically difficult” sequences that stall all optimizers (Supplemental Section “Difficult start sequences”).

### 4.2 Predictive Models and Task Architectures

We benchmark designers on 17 tasks in four categories. The summary of tasks can be found in Table 1 and in the sections below.

#### 4.2.1 Cis-Regulatory Activity (Malinois)

Cis-regulatory elements (CREs) are regions of non-coding DNA which regulate the transcription of neighboring genes. CREs are essential to gene therapies, and to understanding disease mechanisms. We use Malinois models [14] on three different cell-lines, trained on the regulatory output of 700K nucleotide sequences and assayed by MPRAs in a single laboratory.

#### 4.2.2 Transcription Factor Binding (BPNet)

Transcription factors (TFs) are proteins that regulate gene expression by binding to specific DNA sequences. The strength of this binding influences whether genes are activated. Binding affinity is important for understanding gene regulation mechanisms and how genetic variations can disrupt normal cellular function. We use 11 BPNet-lite [12] models from [37] trained on 11 TF datasets from K562 cell-lines in Encyclopedia of DNA Elements (ENCODE) consortium portal [38].

#### 4.2.3 Chromatin Accessibility (BPNet)

Chromatin accessibility is the degree to which DNA is physically available for other molecules to bind to, such as transcription factors. Accessible regions of DNA are more easily targeted by therapies, and DNA access helps understand how a drug might influence gene regulation. We use a BPNet-lite [12] model from [37] trained on ENCODE [38] data.

#### 4.2.4 Mean Ribosomal Loading (RiNALMo)

Optimizing mRNA translational efficiency is crucial for vaccines and protein replacement therapies. The rate of protein synthesis is often determined by ribosome loading on the 5’ UTR. We utilized RiNALMo [13], a 650M-parameter language model pretrained on 36 million RNA sequences, to predict Mean Ribosome Load (MRL). Designers optimized 100-nucleotide 5’ UTR sequences to maximize predicted MRL, testing their ability to navigate the complex syntax of RNA translation regulation.

#### 4.2.5 Cell-type specific gene expression (Enformer)

Selective gene expression is the process where some genes are more active in certain cells, tissues, or at specific times compared to others. It is critical to understanding disease mechanisms, and can be used to predict a patient’s response to treatment. We used the Enformer network [10], accessed through the gRelu library [34], to design sequences that maximized the selectivity of expression. Enformer was trained on thousands of human and mouse DNA sequences using epigenetic and transcriptional datasets, to predict targets such as gene expression and DNA accessibility.

### 4.3 Design Algorithms and Implementation

#### 4.3.1 The Gradient-Guided Adaptive Beam Search (GrAdaBeam) Algorithm

GrAdaBeam (Gradient-guided Adaptive Beam Search) is a hybrid optimization algorithm that unifies evolutionary search with gradient-based guidance (Fig. 3a). For every optimization step, the algorithm decomposes the generation of new candidate sequence *s*^*′*^ in the set of all candidate sequences *S*^*′*^ from a parent *s* into three distinct, probabilistic steps: (1) determining the number of mutations, (2) selecting mutation locations, and (3) selecting the target nucleotides.

##### Number of mutations (Adaptive Mutation Count)

Unlike standard evolutionary algorithms that use a fixed mutation rate or rejection sampling, GrAdaBeam explicitly samples the number of edits *N* for each child from a specialized distribution derived to maximize search efficiency. We reject the null mutation case (*N* = 0) and reallocate its probability mass to ensure every step moves in sequence space:

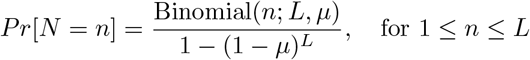

where *L* is the sequence length and *µ* is the adaptive mutation rate hyperparameter. It is determined adaptively using the method described in section “Population Based Training (PBT)” below. This formulation allows for *O*(1) sampling of mutation counts, significantly accelerating the inner loop compared to rejection sampling methods like AdaLead.

##### Mutation location

To determine where to mutate, GrAdaBeam employs a twostage process to balance exploration, exploitation, and computational cost.

###### Stage A: Initial Probability Computation (At Root)

All successful candidates in the beam from the previous round are selected as a “root.” When a sequence is selected as a root, we compute gradient logits *l* via Taylor in silico mutagenesis (TISM) [26]. To ensure stability, logits are standardized by their standard deviation: 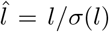. We apply dynamic temperature scaling to prevent the softmax distribution from becoming deterministic. The temperature *τ* is calculated such that the maximum scaled logit does not exceed a threshold *L*_*max*_ (default 3.0):

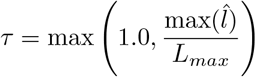

The gradients are converted to probabilities via 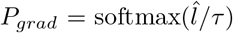. To prevent single dominant positions from collapsing diversity, we clip probabilities at a cap *C* (default 0.10) and re-normalize. Finally, we mix this with a uniform distribution *U* using an exploration coefficient *α*:

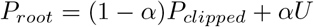

*α* is determined dynamically, as described in the section “Population Based Training (PBT)” below.

###### Stage B: Iterative Masking (During Rollout)

GrAdaBeam does not recompute gradients at every step of a rollout. Instead, it reuses *P*_*root*_. After mutating a set of positions *M*, the probability map for the child node is updated by masking the used locations:

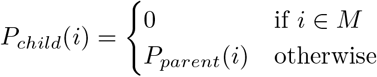

The map is then re-normalized to sum to 1.0. This ensures the rollout sequentially explores the next-best gradient-suggested locations without redundancy or expensive re-computation. This computational shortcut is most applicable in the regime where 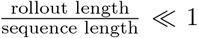 (a rollout is the sequence of consecutive mutation steps from a single root). This shortcut is essential for GrAdaBeam to be time-efficient, given the large relative-compute cost of gradients (Table 1).

###### Selecting the mutation

Once the *n* locations are identified, the specific nucleotide *b*_*new*_ at each location is selected uniformly at random from the set of bases differing from the current base *b*_*old*_. We employ uniform selection here because our ablation studies on gradient-guided methods (Gradient Evo) demonstrated that applying gradient guidance to the value of the edit yielded no statistical performance improvement over random selection (*p* = 0.98), whereas guiding the location was critical.

###### Population Based Training (PBT)

To dynamically tune the search strategy without manual schedules, we employ Population Based Training (PBT). While standard evolutionary algorithms rely on fixed hyperparameters, GrAdaBeam treats the search strategy itself as an evolving entity. Each candidate sequence in the beam carries a tuple of hyperparameters *θ*_*i*_ = *µ, α* . When a high-fitness child sequence is generated and survives the beam cut to become a parent for the next generation, it inherits *θ*_*parent*_. These parameters are then updated using two distinct mechanisms. This allows the algorithm to autonomously transition between global exploration and local refinement as it traverses the fitness landscape.

###### Mutation rate *µ* (standard perturbation)

The mutation rate *µ* governs the number of edits *N* sampled for each step. To adapt this parameter, we employ a standard PBT perturbation mechanism. With probability *p*_*perturb*_ (set to 0.20), the inherited *µ* is multiplied by a factor randomly sampled from {0.8, 1.2}:

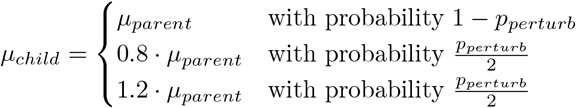

This allows the algorithm to stochastically explore different step sizes, naturally selecting for higher mutation rates in flat regions of the landscape (to escape local optima) and lower rates in steep regions (to fine-tune high-fitness candidates).

###### Exploration alpha *α*

The exploration coefficient *α* balances gradient-based exploitation against uniform exploration. Instead of random perturbation, *α* is updated via a Bayesian inference mechanism based on the *observed* utility of the gradient. For a child sequence with a set of chosen mutation locations *M*, we calculate the posterior probability that these locations were drawn from the uniform distribution *P*_*uniform*_ rather than the gradient distribution *P*_*grad*_:

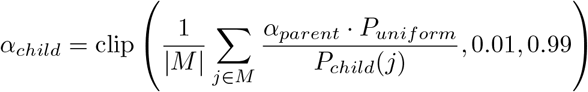

This creates a self-correcting feedback loop: if the algorithm successfully selects mutations that the gradient model deemed unlikely (low *P*_*grad*_), *α* automatically increases, encouraging further exploration. Conversely, if high-gradient locations are selected, *α* decreases to favor exploitation.

###### Computational Efficiency and Scaling

Calculating gradients for large genomic models incurs a significant computational cost compared to forward inference. The cost of a backward pass ranges from 2× (BPNet) to 3.5× (Enformer) that of a forward pass (Table 1). For large models like Enformer, gradient calculations are prohibitively slow for large-scale design. GrAdaBeam addresses this bottleneck through its “Iterative Masking” mechanism, which amortizes the cost of a single gradient calculation over multiple mutation steps (*N*). By performing a rollout of edits using a single cached gradient map, GrAdaBeam reduces the frequency of expensive backward passes. Empirically, this allowed GrAdaBeam to search the Enformer landscape with a wall-clock throughput comparable to inference-only evolutionary methods, while still benefiting from gradient guidance. In contrast, standard gradient methods like Ledidi were constrained by the high latency of the backward pass, completing significantly fewer optimization steps within the fixed time budget.

#### 4.3.2 Baseline Optimization Strategies

##### Baseline algorithm selection

The five baseline designers were chosen to be representative of different types of algorithms, and to reflect what has been used in previous benchmarks. Directed Evolution and AdaLead [33] are common examples of greedy evolutionary algorithms. They are easy to implement, and have been used in previous benchmarks [14, 33–35]. Ledidi [40] is an example of strongly performing, greedy, purely gradient-based approaches. Simulated annealing [39] is the only non-greedy optimizer, and is a common baseline for works in this field [14, 33–35]. We did not include Bayesian optimization [24], CMA-ES [25], or reinforcement learning approaches [23] as baselines, as these require either continuous relaxation of the sequence space, surrogate model fitting, or policy training infrastructure that fundamentally changes the computational setup, making wall-clock-matched comparisons difficult.

##### AdaLead

AdaLead [33] is an adaptive greedy search algorithm based on genetic evolution algorithms. It perturbs the best-known sequences through iterative recombination and mutation. Each round, it makes mutations based on a fixed per-nucleotide probability. It then continues this sequential iteration procedure until the sequence is less fit than the starting sequence, keeping track of every seen sequence along the way and including them in a set of candidates for the next round. When a fixed compute budget is reached, the top percentile candidates in terms of fitness move to the next round, and the process repeats.

##### Ordered and Unordered Beam search

We introduce ordered and Unordered Beam search [27] to nucleic acid design. These designers are staple search algorithms in the computer science literature, and are often used in speech processing [41, 42]. The “beam” in these designers allows them to track multiple strong candidate sequences rather than just the single best sequence. We compare these designers to understand their performance and the impact of fixing edit order. The exact formulations of the two beam algorithms are described in Supplementary Section “Formulation of ordered and unordered beam search”.

#### 4.3.3 Gradient Evo: Hybrid of directed evolution and gradient methods

We introduce a novel designer, gradient-guided Directed Evolution (Gradient Evo), which is a hybrid between the simplicity of Directed Evolution and the performance of Ledidi. The algorithm uses Directed Evolution, where edit locations are guided by Taylor *in silico* mutagenesis (TISM) [26]. TISM approximates the effect of changing nucleotides in a sequence in a computationally efficient way. The equation for TISM, reproduced from [26] is:

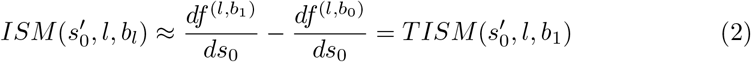

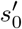:= nucleotide sequence

*b*_0_ := the nucleotide at location *l* in the original sequence

*b*_1_ := the nucleotide at location *l* in the modified sequence

### 4.4 Orthogonal Validation Strategy

To confirm that optimized sequences recover true biological signal rather than exploiting model-specific artifacts, we validated designs using independent “held-out” oracles and motif analysis tools.

#### 4.4.1 Cross-Model Generalization for Ribosomal Loading (Saluki, Optimus 5p)

To validate the mRNA sequences on held-out models, we evaluated the generated 5’ UTR sequences using two independent predictive models: Optimus 5-Prime [15] (predicting mean ribosome load) and Saluki [17] (predicting mRNA half-life). For each generated sequence, we computed the Optimus score using a scanning average of 50bp windows (stride 5) to account for motif positioning, while Saluki scores were computed directly on the full sequence. To assess improvement over the initial seeds, we calculated a “gain” for each sequence by subtracting the score of its corresponding starting sequence. These gains were further normalized to z-scores using the standard deviation of the starting sequences’ scores to allow for comparison across metrics with different scales.

The Ridge Mountain Range Plots (Fig. 4c,d) visualize the distribution of these z-scored gains for each design method, modeled using Kernel Density Estimation (KDE) and ordered vertically by median gain; the red dashed lines indicate the median improvement, illustrating both the central tendency and consistency of each method. The Trade-off Map (Fig. 4b) displays the mean raw scores for translation efficiency (Optimus) versus stability (Saluki), with error bars representing the 95% confidence interval (SEM × 1.96), highlighting the Pareto frontier of multi-objective optimization. Finally, the Performance Heatmap (Fig. 4a) ranks methods by a composite score (sum of standardized Optimus and Saluki scores) and displays the mean standardized performance for each metric, with color intensity normalized column-wise (0–1) to facilitate relative comparison between methods.

#### 4.4.2 Cross-Model Generalization for Gene Expression (Borzoi)

To validate the DNA sequences on held-out models of gene expression, we evaluated the generated sequences using two predictive models: Enformer [10] (the optimization target) and Borzoi [16] (independent validation). Both models were used to predict gene expression maximized for muscle cells, and minimized for liver cells. For Borzoi evaluation, generated sequences were injected into the center of a fixed gene desert background to satisfy the model’s input length requirements. We computed the improvement (“gain”) for each sequence by subtracting the score of its corresponding starting sequence from its optimized score. Raw model outputs were inverted prior to this calculation so that higher values indicate higher predicted expression. These gains were normalized to z-scores using the standard deviation of the start sequences’ scores to allow for comparison across metrics with different distributions.

The Ridge Mountain Range Plots ((Fig. 4f) visualize the distribution of these z-scored Borzoi gains for each design method, modeled using Kernel Density Estimation (KDE) and ordered vertically by median gain. The red dashed lines indicate the median improvement, illustrating the central tendency of each method. The Performance Heatmap (Fig. 4e) ranks methods by a composite score (sum of standardized Enformer and Borzoi scores) and displays the mean standardized performance for each metric, with color intensity normalized column-wise (0–1) to facilitate relative comparison between methods.

#### 4.4.3 Motif Recovery Analysis

To verify that the designed sequences contained the expected regulatory syntax, we utilized the JASPAR 2022 CORE non-redundant vertebrate database [31] as our ground truth. We mapped each target class in the BPNet model to its corresponding JASPAR Position Frequency Matrix (PFM).

##### FIMO Scoring (Fig. 4g)

We quantified motif enrichment using FIMO [29] (Find Individual Motif Occurrences) from the MEME Suite. For each designer-target pair, we scanned both the generated proposal sequences and the initial start sequences for the target’s JASPAR motif. A sequence was considered a “hit” if it contained at least one motif occurrence with a *q*-value *<* 0.1. We calculated the “Delta Hit Rate” as the difference in hit percentages between the proposal and start sets (*HitRate*_*proposal*_ − *HitRate*_*start*_). This metric isolates the contribution of the optimization process, correcting for any motifs present in the initialization. The results were aggregated into a heatmap to compare performance across all experimental conditions.

##### De Novo Motif Discovery (Fig. 4h)

To visually inspect the quality of the generated motifs, we performed de novo motif discovery using STREME [30] (Sensitive, Thorough, Rapid, Enriched Motif Elicitation). For selected representative targets (MYC, ELF4, E2F3), we fed the designed sequences from GrAdaBeam, AdaLead, and Ledidi into STREME to identify enriched patterns without prior knowledge of the target. The resulting discovered motifs were then compared to the JASPAR database using TOMTOM [32] to identify the best match. We visualized the Position Weight Matrices (PWMs) of the best-matching discovered motifs alongside the true JASPAR reference using logomaker, confirming that the algorithms successfully reconstructed the canonical binding preferences of the target TFs.

Matches between de novo discovered motifs and JASPAR references were evaluated using TOMTOM’s Euclidean distance metric. We report the *p*-values for the alignment between the discovered motif and the canonical binding site for the target TF. Matches were considered successful if the discovered motif aligned to the target TF (or its known binding partner, e.g., MAX for MYC) with a TOMTOM *p*-value *<* 0.05. In our representative analysis of MYC, ELF4, and E2F3 (Fig. 4h), all discovered motifs met this criterion with *p*-values significantly below this threshold (range 10^*−*4^ to 10^*−*7^), confirming the identity of the generated regulatory syntax.

### 4.5 Benchmarking Metrics

#### 4.5.1 Optimization Quality

##### Normalized Fitness Gain (Normalized Δ*Z*)

To compare performance across tasks with different dynamic ranges (e.g. log-counts vs. real-valued expressions), we calculate scaled Z-scores of fitness improvements (Fig. 3a). Each designer receives one score per task and start sequence pair. For a start sequence *S*_0_ and designed sequence *S*^*′*^, the raw gain is normalized by the standard deviation of the initial pool (*σ*_*start*_).

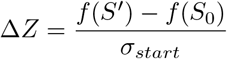

where *f* (*S*) is the fitness of sequence *S* (larger is better), and *σ*_*start*_ is the standard deviation of the fitnesses of the start sequences for the given task.

To facilitate comparison across disparate tasks, these values are then linearly rescaled to the [0, 1] range for each task and start sequence.1.0 represents the maximum gain achieved by any algorithm for that task and start sequence, while 0.0 represents the minimum. The resulting metric allows us to aggregate performance across tasks with vastly different dynamic ranges.

##### Order Score

To quantify the reliability of an algorithm relative to its peers, we calculate the Order Score (Fig. 3b). Each designer receives one order score associated with each task and start sequence pair. For a given designer, task, and start sequence, the order score counts the number of competing algorithms that produced worse final fitnesses. The best possible order score is one less than the number of designers, indicating the top performing algorithm. The worst possible order score is zero, indicating that all other algorithms performed better.

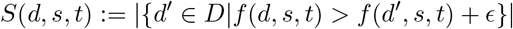

where *d* is the designer; *s* is the start sequence; *t* is the task; *D* is the set of all designers; *f* (*d, s, t*) is the final fitness with designer *d* (larger is better), start sequence *s*, and task *t*; and *ϵ* is a small margin to avoid small differences from affecting the score (*ϵ* = 0.001 in our study).

The order score allows a fair comparison across tasks that have different numerical ranges and different distributions of fitness scores. It is somewhat preferable to scores that take magnitude into account because the large number of order scores allows us to measure designers that consistently outperform others (the number of order scores for each designer in Fig. 3b is 1,600).

#### 4.5.2 Sequence Diversity

To measure mode collapse, we compute the average pairwise Hamming distances between final sequences (Fig. 3c). Each designer receives one sequence diversity score per task.

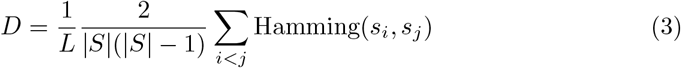

where *S* is the set of final sequences generated by the algorithm for a given task (aggregated across all start sequences), *L* is the number of editable nucleotides (sequence length), and Hamming(*s*_*i*_, *s*_*j*_) counts the number of positions where the nucleotides differ. This metric captures the variability of the candidate pool, distinguishing algorithms that converge to a single mode from those that explore diverse local minima. Higher diversity indicates the algorithm has found multiple distinct local optima, whereas low diversity suggests collapse to a single consensus sequence. For Fig. 3c, this diversity rate is standardized to a Z-score for each task (inverted so that higher diversity yields a higher score) and then min-max normalized to [0, 1].

#### 4.5.3 Stability: Start Sequence & Random Seed

We quantify algorithmic stability in two dimensions ((Fig. 3c):

- **Start Sequence Stability**: We quantify the robustness of an algorithm to the choice of start sequence. For each task and algorithm, we computed the standard deviation of the mean final energy achieved across all 100 distinct start sequences. Lower variance indicates less sensitivity. This aggregate metric was log-transformed (log(1+*x*)) and standardized (Z-scored) relative to the distribution of stability scores of all algorithms for that task. The Z-scores were inverted and min-max normalized to the [0, 1] interval, with bounds defined by the minimum and maximum scores across all algorithms within the task.
- **Random Seed Stability**: We quantify the robustness of an algorithm to stochas-ticity during optimization. For each task, algorithm, and start sequence, we calculated the standard deviation of the final energy values across *n* ≥ 5 independent random seeds. Lower variance indicates a more reliable algorithm. To account for the heavy-tailed distribution of variances, we applied a log-transformation (log(1 + *x*)). These values were then standardized (Z-scored) relative to the distribution of log-variances across all algorithms and start sequences within the same task. Finally, to facilitate comparison, the Z-scores were inverted (as lower variance indicates higher stability) and min-max normalized to the [0, 1] interval, where the normalization bounds were defined by the minimum and maximum scores of all algorithms for that specific start sequence.

### 4.6 Statistical Analysis of In-Silico Performances

To assess the statistical significance of the performance differences between GrAdaBeam and the baseline algorithms, we employed a rigorous non-parametric testing framework. For each optimization task, algorithms were evaluated on 100 paired independent runs (matched by start sequence). Tasks where all algorithms converged to indistinguishable performance levels (defined as a spread in mean energy of less than 0.5 units) were excluded from the statistical analysis to avoid ceiling effects.

We performed pairwise comparisons using the Wilcoxon signed-rank test [43]. For task-level analysis, we compared the distribution of final energy values between GrAdaBeam and each baseline across the 100 paired runs. For the global analysis across the entire benchmark suite, we first computed the mean rank of each algorithm for each task, and then applied the Wilcoxon signed-rank test to the paired distributions of these task-level mean ranks. To control for the family-wise error rate across multiple pairwise comparisons, all *p*-values were adjusted using the Holm-Bonferroni correction [44], with statistical significance determined at the *α* = 0.05 level.

### 4.7 Quantifying the performance divide between gradient and evolutionary methods

To evaluate the domain-specific advantages of different optimization paradigms, we stratified the algorithms into two categories: evolutionary methods and gradient-based methods (Table 2, excluding GrAdaBeam). For every task, we determined the representative performance of each category by selecting the algorithm that achieved the best mean metric. We quantified the comparative advantage using the normalized relative difference, calculated as 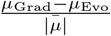, where *µ*_Grad_ and *µ*_Evo_ correspond to the mean scores of the top-performing evolutionary and gradient algorithms, respectively, and 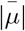 is the mean of the absolute values of the two top-performing mean scores.

**Table 2.** Design algorithms benchmarked in NucleoBench, with classification by optimization paradigm. Above the solid line are designers already found in the nucleic acid design literature. Below the line are designers from the search literature not previously used to benchmark nucleic acid sequence design and hybrid algorithms devised in this work.

| Algo | Description | Gradient-based |
| --- | --- | --- |
| Directed Evolution | Random mutations, track the best. | ✗ |
| Simulated Annealing [39] | Greedy optimization with random jumps. | ✗ |
| AdaLead [33] | Heuristic-guided random-mutation search. | ✗ |
| Ledidi [40] | Gumbel softmax sampling for maximal input. | ✓ |
| Ordered Beam [27] | Greedy beam search, in fixed edit order. | ✗ |
| Unordered Beam [27] | Greedy beam search, any edit order. | ✗ |
| Gradient Evo | Directed Evolution, guided by gradients. | ✓ |
| GrAdaBeam | Efficient beam search guided by gradients. | ✓ |

Uncertainty in the relative difference was estimated by propagating the standard errors of the means (*σ*_Evo_, *σ*_Grad_) under the assumption of independence, yielding a combined standard error for the normalized difference. We computed 95% confidence intervals as ±1.96 times the combined standard error. Differences were classified as statistically significant if the confidence interval excluded zero.

## Supplementary information

The online version contains supplementary material available at [insert DOI/URL once accepted].

## Acknowledgements

We thank Sager Gosai for his invaluable guidance on interpreting motifs and task selection. We thank Daniel Friedman for early discussions on formalizing edit order. We thank Anna Lewis and Vikram Agarwal for guidance throughout the paper writing process.

## Declarations

### Ethics approval and consent to participate

N/A

### Consent for publication

The authors consent to publish

### Data availability

https://zenodo.org/records/15775704

### Code availability

https://github.com/move37-labs/nucleobench

### Author contributions

J.S. conceived and designed the study. J.S. performed data acquisition and ingestion. J.S, E.S. performed algorithm development. J.S., C.Y.M. performed experiments and analyzed results. J.S. wrote the initial version of the paper. All authors had access to the data and have edited, reviewed and approved the final version of the paper for publication. All authors accept responsibility for the accuracy and integrity of all aspects of the research.

