## Supplemental Information for "GrAdaBeam: Combining model gradients with evolutionary search for generalizable nucleic acid design"

### Supplementary Information

#### 1 Per-task performance table

**Table 1**: 95% confidence intervals for the best per-task results. For each (task, algorithm, hyperparameters), the 95% confidence interval was computed across start sequences and random seeds using the final optimized energies (lower is better). Red is the best, blue second best, per column.

| Algorithm | CRE | | | TF binding | | | | | | | | | | | ATAC | Enformer |
|---|---|---|---|---|---|---|---|---|---|---|---|---|---|---|---|---|
| | K562 | HepG2 | SK-N-S | CTCF | E2F3 | ELF4 | GATA2 | JUNB | MAX | MECOM | MYC | OTX1 | RAD21 | SOX6 | | ↑muscle ↓liver |
| Directed Evolution | **-15.4, -15.3** | -14.5, -14.2 | -14.2, -13.8 | -0.9, -0.7 | -1.4, -1.3 | 0.6, 0.7 | -0.9, -0.7 | -4.1, -4.1 | -0.2, -0.0 | -1.6, -1.3 | -0.1, 0.0 | -4.5, -4.4 | -2.1, -1.9 | -3.4, -3.3 | -2.2, -1.9 | -10319, -5750 |
| Simulated Annealing | **-15.4, -15.4** | -10.4, -10.3 | -8.2, -8.2 | -3.7, -3.7 | -2.8, -2.8 | -2.3, -2.3 | -3.0, -2.9 | -4.5, -4.5 | -2.7, -2.7 | -2.5, -2.4 | -1.4, -1.4 | -4.8, -4.8 | -4.0, -4.0 | -4.1, -4.1 | -3.8, -3.8 | -12682, -6788 |
| AdaLead | **-15.4, -15.4** | **-15.4, -15.4** | **-15.4, -15.4** | -14.9, -14.7 | -25.2, -24.9 | -18.5, -18.2 | -19.4, -18.9 | **-17.3, -17.3** | -17.4, -17.2 | -36.7, -36.1 | -15.7, -15.3 | **-5.9, -5.9** | -17.3, -16.8 | -31.7, -31.1 | **-38.3, -37.2** | -18919, -11011 |
| Ledidi | **-15.4, -15.4** | **-15.4, -15.4** | **-15.4, -15.3** | **-18.2, -17.7** | **-28.6, -28.4** | **-20.8, -20.5** | **-27.0, -26.4** | -15.7, -15.4 | **-20.6, -20.3** | **-40.3, -39.7** | **-16.4, -15.9** | -4.6, -4.6 | **-25.3, -24.5** | **-33.0, -32.7** | **-46.8, -45.8** | **-1186192, -1040939** |
| Ordered Beam | **-15.4, -15.4** | **-15.4, -15.4** | **-15.4, -15.3** | -6.1, -6.0 | -10.0, -9.9 | -6.9, -6.8 | -8.6, -8.4 | -7.3, -7.2 | -7.5, -7.3 | -11.8, -11.6 | -6.5, -6.3 | -5.3, -5.2 | -7.5, -7.3 | -12.0, -11.8 | -2.4, -2.1 | -14972, -8515 |
| Unordered Beam | **-15.4, -15.4** | **-15.4, -15.4** | **-15.4, -15.4** | -14.0, -13.6 | -22.7, -22.4 | -17.0, -16.9 | -16.8, -16.6 | -16.9, -16.8 | -14.9, -14.6 | -33.0, -32.4 | -13.8, -13.4 | **-5.8, -5.8** | -15.1, -14.2 | -29.3, -28.9 | -31.4, -30.8 | -21869, -12796 |
| Gradient Evo | **-15.4, -15.4** | **-15.2, -14.9** | -14.6, -13.9 | -14.4, -14.0 | -23.1, -22.8 | -17.1, -16.7 | -17.1, -16.8 | -17.0, -16.9 | -15.3, -15.0 | -33.3, -32.6 | -13.7, -13.3 | **-5.8, -5.8** | -15.4, -14.5 | -29.8, -29.2 | -35.7, -34.7 | -11130, -5962 |
| AdaBeam | **-15.4, -15.4** | **-15.4, -15.4** | **-15.4, -15.4** | **-17.2, -16.9** | **-32.1, -32.0** | **-23.4, -23.0** | **-25.7, -25.1** | **-17.6, -17.5** | **-19.7, -19.4** | **-43.7, -43.3** | **-22.9, -22.4** | **-5.9, -5.9** | **-22.5, -21.5** | **-36.4, -36.0** | **-46.9, -46.1** | -23894, -13523 |

The full list of per-task performances are in table 1.

#### 2 Per-task variability due to random seed

The full list of per-task performance variability due to random seed are in table 2.

#### 3 Per-task variability due to start sequence

The full list of per-task performance variability due to start sequences are in table 3.

#### 4 Difficult start sequences

An outstanding question is how correlated performance fluctuations are with the start sequence, and whether there exist "intrinsically difficult start sequences." To that end, we analyzed whether certain sequences are likely to be responsible for poor performance in an optimization-agnostic way. For each task, we look at the experiment results from the best hyperparameters for each algorithm. We perform the non-parametric Friedman Test and the Nemenyi posthoc test by treating the 100 start sequence identities as nominal variables, the different designers as treatments, and the performances as continuous Y variables.

**Table 2**: Performance variability based on random seed. For each (task, algorithm, hyperparameters), we show the (25th percentile, 75th percentile) for the best performing hyperparameters.

| Algorithm | CRE | | | TF binding | | | | | | | | | | | ATAC | Expression |
|---|---|---|---|---|---|---|---|---|---|---|---|---|---|---|---|---|
| | K562 | HepG2 | SK-N-S | CTCF | E2F3 | ELF4 | GATA2 | JUNB | MAX | MECOM | MYC | OTX1 | RAD21 | SOX6 | | $\uparrow$muscle $\downarrow$liver |
| Directed Evolution | (0.00, 0.00) | (0.95, 1.97) | (1.32, 2.35) | (0.35, 0.78) | (0.12, 0.29) | (0.15, 0.42) | (0.41, 0.72) | (0.06, 0.13) | (0.28, 0.62) | (0.11, 0.32) | (0.30, 0.55) | (0.01, 0.03) | (0.36, 0.72) | (0.09, 0.21) | (0.36, 0.73) | (646.21, 3273.90) |
| Simulated Annealing | (0.00, 0.00) | (0.26, 0.49) | (0.18, 0.28) | (0.08, 0.12) | (0.07, 0.12) | (0.04, 0.06) | (0.08, 0.15) | (0.01, 0.02) | (0.07, 0.11) | (0.05, 0.09) | (0.06, 0.10) | (0.00, 0.01) | (0.09, 0.14) | (0.05, 0.08) | (0.06, 0.10) | (694.23, 6233.92) |
| AdaLead | (0.00, 0.00) | (0.00, 0.00) | (0.00, 0.05) | (0.44, 0.75) | (0.45, 0.77) | (0.42, 0.76) | (0.56, 0.87) | (0.10, 0.15) | (0.33, 0.58) | (1.00, 1.63) | (0.59, 0.88) | (0.01, 0.01) | (0.56, 1.07) | (0.87, 1.45) | (1.38, 2.11) | (1051.72, 7289.75) |
| Ledidi | (0.00, 0.00) | (0.00, 0.01) | (0.00, 0.02) | (0.56, 1.09) | (0.42, 0.68) | (0.43, 0.67) | (0.70, 1.15) | (0.45, 0.74) | (0.35, 0.67) | (0.93, 1.53) | (0.67, 1.07) | (0.01, 0.02) | (0.90, 1.48) | (0.46, 0.82) | (1.53, 2.43) | (120364.95, 243801.88) |
| Ordered Beam | (0.00, 0.00) | (1.50, 1.88) | (1.86, 2.24) | (0.36, 0.54) | (3.97, 4.74) | (2.69, 3.24) | (1.11, 1.91) | (1.62, 2.23) | (0.94, 1.53) | (7.15, 8.38) | (2.14, 3.52) | (0.03, 0.05) | (1.32, 2.02) | (4.70, 5.63) | (0.29, 0.52) | (778.27, 6819.78) |
| Unordered Beam | (0.00, 0.00) | (0.00, 1.54) | (1.05, 2.19) | (0.50, 0.82) | (0.47, 0.85) | (0.46, 0.71) | (0.57, 0.83) | (0.12, 0.19) | (0.50, 0.75) | (1.11, 1.66) | (0.55, 0.91) | (0.01, 0.01) | (0.65, 1.71) | (1.03, 1.66) | (1.73, 2.77) | (931.13, 7811.52) |
| Gradient Evo | (0.00, 0.00) | (1.14, 1.86) | (1.66, 2.51) | (0.53, 0.83) | (0.60, 0.82) | (0.54, 0.75) | (0.52, 0.81) | (0.14, 0.20) | (0.44, 0.72) | (1.26, 1.88) | (0.60, 0.87) | (0.01, 0.01) | (0.74, 1.35) | (1.13, 1.77) | (1.63, 2.62) | (623.83, 2415.36) |
| AdaBeam | (0.00, 0.00) | (0.00, 0.04) | (0.54, 1.08) | (0.44, 0.71) | (0.35, 0.53) | (0.39, 0.70) | (0.96, 1.47) | (0.08, 0.14) | (0.32, 0.57) | (0.74, 1.18) | (0.58, 0.99) | (0.01, 0.01) | (0.84, 1.40) | (0.64, 1.01) | (1.29, 1.91) | (1157.73, 6996.59) |

**Table 3**: Performance variability based on start sequences. For each (task, algorithm, hyperparameters), We show the (mean +- standard deviation) for the best performing hyperparameters.

| Algorithm | CRE | | | TF binding | | | | | | | | | | | ATAC | Expression |
|---|---|---|---|---|---|---|---|---|---|---|---|---|---|---|---|---|
| | K562 | HepG2 | SK-N-S | CTCF | E2F3 | ELF4 | GATA2 | JUNB | MAX | MECOM | MYC | OTX1 | RAD21 | SOX6 | | $\uparrow$muscle $\downarrow$liver |
| Directed Evolution | -15.32 ± 0.19 | -14.36 ± 0.85 | -13.99 ± 0.89 | -0.83 ± 0.51 | -1.37 ± 0.19 | 0.65 ± 0.43 | -0.79 ± 0.39 | -4.07 ± 0.07 | -0.13 ± 0.53 | -1.44 ± 0.57 | -0.04 ± 0.28 | -4.44 ± 0.04 | -1.99 ± 0.57 | -3.34 ± 0.11 | -2.02 ± 0.66 | -8035.22 ± 11514.42 |
| Simulated Annealing | -15.39 ± 0.00 | -10.33 ± 0.19 | -8.18 ± 0.11 | -3.66 ± 0.05 | -2.78 ± 0.05 | -2.30 ± 0.02 | -2.96 ± 0.06 | -4.49 ± 0.01 | -2.69 ± 0.04 | -2.45 ± 0.03 | -1.38 ± 0.04 | -4.77 ± 0.00 | -4.03 ± 0.05 | -4.12 ± 0.03 | -3.82 ± 0.04 | -9735.61 ± 14852.95 |
| AdaLead | -15.39 ± 0.00 | -15.39 ± 0.00 | -15.39 ± 0.00 | -14.79 ± 0.66 | -25.03 ± 0.71 | -18.35 ± 0.84 | -19.12 ± 1.15 | -17.30 ± 0.13 | -17.29 ± 0.61 | -36.43 ± 1.46 | -15.48 ± 0.86 | -5.87 ± 0.01 | -17.04 ± 1.39 | -31.42 ± 1.50 | -37.74 ± 2.58 | -14965.50 ± 19927.95 |
| Ledidi | -15.39 ± 0.00 | -15.38 ± 0.02 | -15.36 ± 0.07 | -17.93 ± 1.16 | -28.50 ± 0.57 | -20.64 ± 0.72 | -26.68 ± 1.60 | -15.55 ± 0.87 | -20.47 ± 0.67 | -40.01 ± 1.51 | -16.14 ± 1.22 | -4.64 ± 0.03 | -24.87 ± 2.02 | -32.84 ± 0.74 | -46.32 ± 2.55 | -1113565.75 ± 366020.88 |
| Ordered Beam | -15.39 ± 0.00 | -15.39 ± 0.00 | -15.34 ± 0.26 | -6.01 ± 0.25 | -9.93 ± 0.36 | -6.88 ± 0.26 | -8.52 ± 0.41 | -7.21 ± 0.27 | -7.39 ± 0.43 | -11.72 ± 0.59 | -6.39 ± 0.48 | -5.25 ± 0.02 | -7.38 ± 0.61 | -11.93 ± 0.57 | -2.22 ± 0.67 | -11743.73 ± 16270.57 |
| Unordered Beam | -15.39 ± 0.00 | -15.39 ± 0.00 | -15.39 ± 0.00 | -13.79 ± 0.87 | -22.53 ± 0.83 | -16.95 ± 0.47 | -16.71 ± 0.49 | -16.85 ± 0.17 | -14.79 ± 0.79 | -32.71 ± 1.67 | -13.62 ± 0.96 | -5.83 ± 0.01 | -14.62 ± 2.22 | -29.13 ± 0.92 | -31.11 ± 1.66 | -17332.86 ± 22861.82 |
| Gradient Evo | -15.39 ± 0.00 | -15.06 ± 0.96 | -14.26 ± 1.95 | -14.19 ± 0.87 | -22.96 ± 0.75 | -16.89 ± 0.84 | -16.98 ± 0.82 | -16.96 ± 0.17 | -15.13 ± 0.85 | -32.91 ± 1.72 | -13.47 ± 0.90 | -5.84 ± 0.01 | -14.95 ± 2.50 | -29.48 ± 1.48 | -35.17 ± 2.49 | -8546.06 ± 13022.21 |
| AdaBeam | -15.39 ± 0.00 | -15.39 ± 0.00 | -15.39 ± 0.00 | -16.96 ± 0.77 | -31.74 ± 0.54 | -23.03 ± 0.66 | -24.92 ± 1.73 | -17.53 ± 0.13 | -18.21 ± 1.19 | -41.19 ± 1.15 | -20.92 ± 1.00 | -5.89 ± 0.01 | -21.24 ± 2.80 | -35.97 ± 1.09 | -40.95 ± 8.24 | -18708.53 ± 26135.83 |

Table 4 shows that intrinsically difficult start sequences exist, and are not uniformly distributed between tasks. Specifically, we observed roughly four categories of problem: no difficult seeds (8/16), 1 difficult seed (6/16), more than 1 and fewer than 5 difficult seeds (1/16), and many ($\geq$ 5%) difficult seeds (1/16). Cell-specific expression tasks and the majority of transcription factor binding tasks did not have any hard sequences.

**Table 4**: Top row) Friedman test for the existence of extremal start sequences, then the Nemenyi post hoc test for identifying specific start sequences. Task name in the top row. Middle row is (fraction of seeds that are extremal, Friedman test statistic, significance). Bottom row) Aggregate amount of time for optimization convergence. Convergence times were computed for each task / optimizer, then converted to a z-score across tasks per optimizer. Z-scores were then averaged for each task. Higher scores indicate that this task took optimizers longer than others to converge. See "Performance variability from start sequence" in Methods for a description of the order score.

| **Task type** | | **CRE** | | | **TF binding** | | | | | | | | | | | **ATAC** | **Expression** |
|---|---|---|---|---|---|---|---|---|---|---|---|---|---|---|---|---|---|
| | | **K562** | **HepG2** | **SK-N-SH** | **CTCF** | **E2F3** | **ELF4** | **GATA2** | **JUNB** | **MAX** | **MECOM** | **MYC** | **OTX1** | **RAD21** | **SOX6** | | **DNase (↑ muscle ↓ liver)** |
| Existence of difficult start sequences | Fraction of extremal seeds (out of 100) | - | - | - | - | - | 1 | 1 | - | 3 | 1 | - | 1 | 1 | - | 1 | 54 |
| | Friedman test statistic | - | - | - | - | - | 150 | 167 | - | 204 | 141 | - | - | 206 | - | 154 | 447 |
| | Friedman test significance | - | - | - | - | - | 7.6 E-4 | 2.4 E-5 | - | 3.0 E-9 | 3.4 E-3 | - | - | 1.8 E-9 | - | 3.4 E-4 | 1.2 E-45 |
| Optimizer convergence | Order ↓ | 9.4 | 7.4 | 5.3 | 5.7 | 7.7 | 5.0 | 5.0 | 8.6 | 5.7 | 4.1 | 6.7 | 12.6 | 5.9 | 7.3 | 8.7 | 17.5 |

### 5 Number of hyperparameters

**Table 5**: Number of hyperparameters that each designer was evaluated on for the full NucleoBench. Hyperparameters were chosen from the results of smaller experiments.

| **Designer** | **Number of hyperparameters** |
|---|---|
| Directed Evolution | 1 |
| Simulated Annealing | 6 |
| AdaLead | 98 |
| Ledidi | 10 |
| Ordered Beam | 8 |
| Unordered Beam | 34 |
| Gradient Evo | 21 |
| GrAdaBeam | N |

Table 5 has the number of hyperparameters that each designer was evaluated on for the full NucleoBench. Hyperparameters were chosen from the literature, as well as the results of numerous smaller-scale experiments. There were roughly an equal number of hyperparameters tried for each designer that were evaluated on a subset of NucleoBench.

### 6 Formulation of ordered and unordered beam search

For both beam search designers, we decompose edit selection into two independent steps: selecting which location to edit, then selecting what the edit should be.

$$\begin{aligned}\Pr[\text{bp at location } n_t \text{ is changed to } m|H_{t-1}] = \\ \Pr[m|\text{edit location } n_t]\cdot \\ \Pr[\text{edit location } n_t|H_{t-1}]\end{aligned}$$

where $H_{t-1}$ is the relevant history of edits that the algorithm has made for the first $t-1$ edits. For Ordered Beam, we select a random permutation of edit order at the start of the algorithm, and proceed to make edits in that order. The designer stops when all locations have been edited once. Note that this allows any start sequence to be edited to any other sequence. Thus, if $l_i$ is our predetermined edit order, then for Ordered Beam, we have:

$$\Pr[\text{edit location } n_t|H_{t-1}] = \begin{cases} 1 & \text{if } n_t = l_t \\ 0 & \text{otherwise} \end{cases}$$

For Unordered Beam search, we select a single location uniformly at random to edit. Thus, for Unordered Beam:

$$\begin{aligned}\Pr[\text{edit location } n_t|H_{t-1}] &= P(\text{edit location } n_t) \\ &= 1/\text{sequence length}\end{aligned}$$

For both algorithms, the new nucleotide is selected uniformly at random to ensure that it is modified.
